# Social networks in the presence and absence of visual cues

**DOI:** 10.1101/432336

**Authors:** David Bierbach, Stefan Krause, Pawel Romanczuk, Juliane Lukas, Lenin Arias-Rodriguez, Jens Krause

**Affiliations:** Department of the Biology and Ecology of Fishes, Leibniz-Institute of Freshwater, Ecology and Inland Fisheries, Berlin 12587, Germany; Department of Electrical Engineering and Computer Science, Lübeck University of Applied Sciences, Lübeck 23562, Germany; Department of Biology, Institute for Theoretical Biology, Humboldt-Universität zu Berlin, Berlin, Germany; Faculty of Life Sciences, Humboldt-Universität zu Berlin, Thaer Institute, Invalidenstrasse 42, 10115 Berlin, Germany; Bernstein Center for Computational Neuroscience, Humboldt Universität zu Berlin, Philippstr 13, 10115 Berlin, Germany; División Académica de Ciencias Biológicas, Universidad Juárez Autónoma de Tabasco (UJAT), C.P. 86150 Villahermosa, Tabasco, México

**Keywords:** Markov Chain, Social Network Analysis, Poecilia, cave fish, social

## Abstract

We compared the social dynamics of two populations of the live-bearing Atlantic molly (*Poecilia mexicana*) that live in adjacent habitats with very different predator regimes: cave mollies that inhabit a low-predation environment inside a sulfidic cave with a low density of predatory water bugs (*Belostoma* sp.), and mollies that live directly outside the cave (henceforth called “surface” mollies) in a high-predation environment with a high density of fish-eating birds. We filmed the social interactions of marked fish in both environments and analysed their social network dynamics using a Markov model under two different fish densities of 12 and 6 fish per 0.36 m^*2*^. As expected, surface mollies spent overall much more time social than cave mollies. This difference in overall social time was a result of surface mollies being less likely to discontinue social contact (once they had a social partner) and being more likely to resume social contact (once alone) than cave mollies. Interestingly surface mollies were also less likely to leave a current social partner than cave mollies. At low density, mollies (in both environments) were expected to show reduced social encounters which should dramatically change their social dynamics. Surface mollies, however, displayed an ability to maintain their social dynamics at low density (primarily by reducing the convex polygon spanned by the group) which was not observed in cave mollies. Despite the fact that we only compared two populations, our data provide a mechanistic explanation for density compensations of social dynamics that have also been observed in other fish species and give an example of how comparisons between the social dynamics of different populations can be made that go beyond conventional network analyses.

## Introduction

In many group-living species we observe that individuals spent some of their time alone and keep switching between social and solitary periods [1]. This type of social dynamics is widespread and has received particular attention in studies on fish. Anybody who has ever watched fish in an aquarium will have noticed the extremely frequent fission-fusion dynamics at the rate of a few seconds that many species display. Recent studies on the shoaling dynamics of guppies (*Poecilia reticulata*, [2]) as well as juvenile lemon sharks (*Negaprion brevirostris*, [3]) in the wild showed that the association duration between individuals follows a geometric distribution (i.e. with short associations being very frequent and long ones very rare). It has been argued that a geometric distribution could be a good shoaling strategy in response to constant predator threats: if a predator is almost always present and monitors prey for moments when it is alone (e.g., the pike cichlid *Crenicichla frenata* which is the main guppy predator in Trinidad; [4, 5]), then a social dynamic that follows a geometric distribution could be adaptive because there would be no typical period for which the predator has to wait for the prey to be alone (see [2] for a discussion). This view is supported by the fact that fish try to compensate changes in environmental conditions like group density, or water depth by somehow actively maintaining a certain social dynamic: Even when observed over several years and when translocated to different habitats, social dynamics remained largely unchanged in the guppy [2, 6]. Furthermore, it is known that the intensity of perceived predation risk can affect the temporal aspects of social dynamics. Social associations among guppies from high predation populations were found to last longer than those among fish in low predation populations [7]. However, to date no mechanism for the maintenance of social dynamics has been proposed and thus an in-depth analysis on how selection might have operated to shape the compensatory abilities is still lacking.

Here we compared the social network structures and social dynamics in two populations of the same species, the livebearing Atlantic molly (*Poecilia mexicana*) that differ in their evolutionary history as well as in their actual habitat. In the South of Mexico, ancestral forms of the Atlantic molly (in the following referred to as ‘molly’) colonized sulfidic springs as well as cave environments about 100’000 years ago [8–10]. Cave-dwelling mollies, that still possess functional albeit smaller eyes, were found to be less social compared to their surface-dwelling counterparts [11–13] that live only a few meters away outside the cave and share the same H_2_S-containing water [14]. The lower sociality was in part attributed to the cave environment which is free of piscivorous fishes as well as birds, thus predators that hunt using visual cues [11, 15, 16]. As a result, it is assumed that cave mollies do not experience a strong selection pressure to maintain shoaling as an anti-predator behaviour [13].

As a basis for the investigation of the social dynamics of the mollies we used the Markov chain model of Wilson et al. [6], which has been applied to the social systems of the closely related guppy [2, 6, 17]. It describes the social behaviour common to all focal individuals as sequences of ‘behavioural states’ and provides the probabilities of fish switching between these states (see methods for more details). In contrast to previous studies on shoaling behaviour which usually looked only at percentage time spent being social, the above approach allows us to decompose the social dynamics into separate behavioural components which potentially offers a more detailed insight into how selection has acted on fish behaviour. We can assess the probability of fish to stay alone once they are alone and to stay social once they are social. In addition, we can also look at the probability of switching social partners within the period of social time. It has been postulated, for example, that fish from high-predation populations should engage in less partner switching and instead invest in stronger social links with fewer individuals to facilitate cooperative anti-predator behaviours [7, 18, 19].

Specifically, we predicted that individuals in the surface population which is exposed to higher levels of predation (see [20]) should show (i) a lower probability of ending social contact, (ii) a higher probability to join a new social partner once they are alone and (iii) a lower probability of leaving a current nearest neighbour and of switching social partners. We also predicted that the surface-dwelling mollies should be able to maintain their social dynamics despite changes in fish density (as has been found in other fish species, see above). However, for the cave-dwelling population it should be hard to do so in the absence of visual cues although short-ranged lateral line cues might be available for them [21]. Therefore we predicted that the cave mollies should largely follow a spatial null-model in their movements.

## Methods

### Study system

Our study sites are located near the Mexican city of Tapijulapa (Fig 1, Tabasco, United Mexican States). Here, the Río Tacotalpa and its tributaries drain through the mountains of the Sierra Madre de Chiapas and several sulphide spring complexes of volcanic origin have been discovered in the foothills of the Sierra Madre. This area is also rich on natural cave formations, two of which are known to harbour endemic populations of the otherwise widely distributed Atlantic molly, *Poecilia mexicana* (the so-called ‘cave molly’; see [21–23]). The Cueva del Azufre (aka Cueva del Sardinia or Cueva Villa Luz) is divided into 13 different chambers, with Chamber XIII being the innermost chamber (after [22], see map of the cave in Fig 1). The front chambers obtain some dim light through natural skylights in the cave’s ceiling, whereas the rearmost cave chambers (from Chamber VI onwards) lay in complete darkness. Several springs in the cave (mainly in Chamber X) release sulfidic water, and the creek that flows through the cave eventually leaves the underground and turns into the sulfidic El Azufre River. H_2_S concentrations in the cave as well as in the EL Azufre river are high (23µM to 320µM,) and accompanied with very low oxygen levels (<1.5 mg/L, see [14])

**Fig 1:**
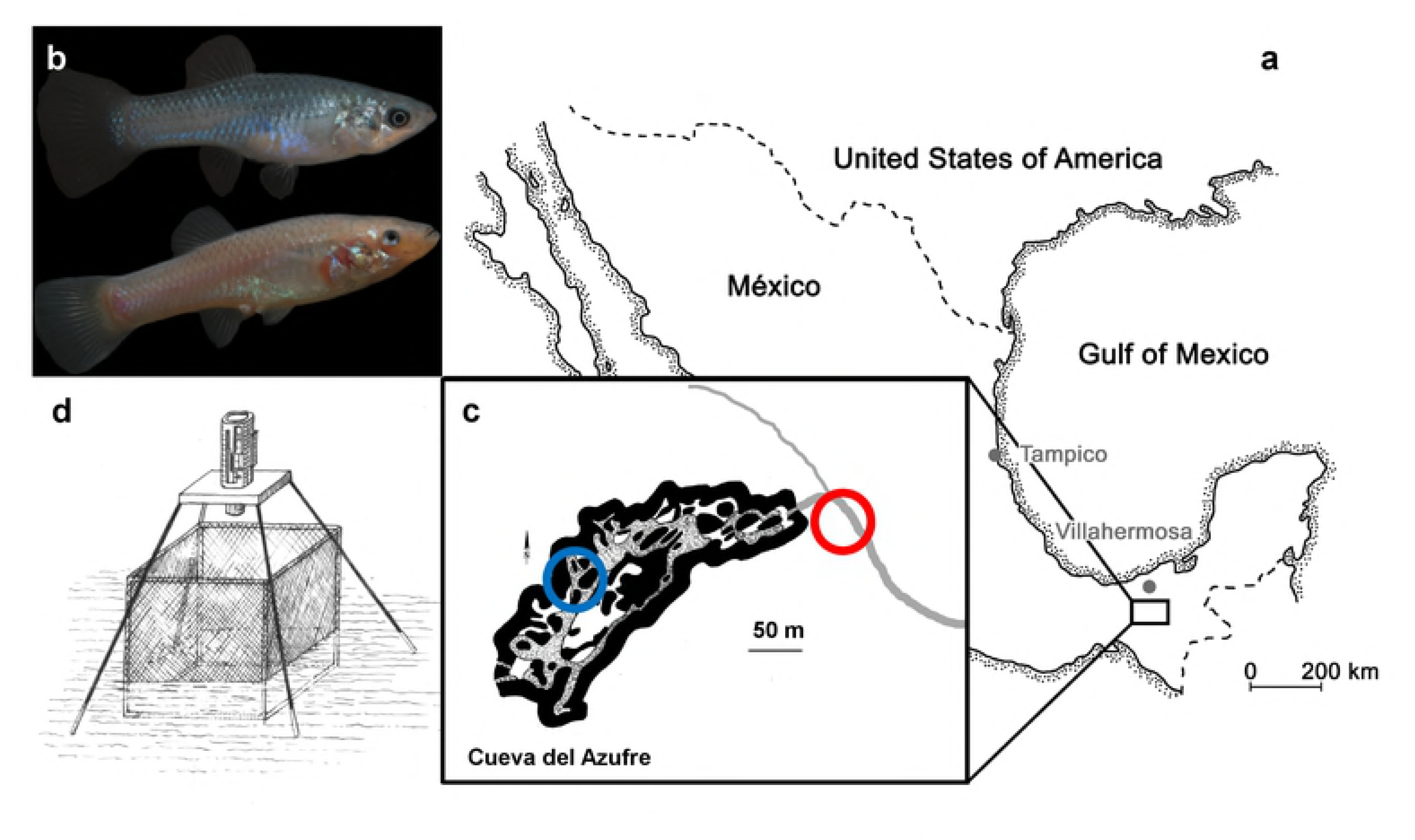
The study system. (a) Both tested molly populations originate from the South of Mexico near the city of Tapijulapa, federal state of Tabasco. Here, ancestral forms of *Poecilia mexicana* colonized both surface (b top, picture of a surface-dwelling molly) as well as cave (b bottom, picture of cave molly) habitats. (c) Locations of the study sites are indicated with red (cave) and blue (surface) circles along the El Azufre river that flows from right to left, i.e., out of the cave. On site, individually marked fish were transferred into a net cage and recorded with an infrared camera (d).

The surface form of the Atlantic molly is widespread in freshwaters along the Central American Atlantic coast [24]. This livebearing species from the family Poeciliidae relies on internal fertilization (males transfer sperm via an adapted anal fin, the Gonopodium, see [25]) and, in contrast to some other molly species in which large males show elaborate courtship displays [26, 27], even large-bodied Atlantic molly males do not court and defend distinct territories [28]. They mostly rely on consensual mating, as females have a mating preference for large male body size [29], while small males often exhibit “ambushing” behaviour (i.e., they hide near groups of females and attempt forced copulations [30]). Nevertheless, males form dominance hierarchies by means of aggressive combat [31, 32]. Although aggressive behaviour is reduced in the cave molly [21, 31–33], males still perform all kinds of aggressive as well as sexual behaviours [28]. In addition, cave mollies have reduced but still functional eyes, which are, however, slightly smaller than those of surface-dwelling mollies [34–37].

### Experimental setup

#### Overview

Our aim was to compare the dynamics of social networks formed by sulphide-adapted surface mollies (in light) with those of sulphide-adapted cave mollies (in total darkness). To do so, we caught and marked fish from two sites located either in the El Azufre river or in cave chamber VII (see Fig 1b) and introduced small groups (N=12, equal sex ratio) of fish into a custom made net cage that was placed at the respective site (Fig 1c). After one day of acclimation, we filmed these groups with a camera under infrared light as *P. mexicana* does not possess infrared-sensitive photoreceptors [37]. To see how social dynamics change with altered fish density, after a first recording 6 fish were haphazardly removed from the cage and the remaining 6 (equal sex ratio) were videotaped again. Afterwards, the removed 6 individuals were put back and the group of 12 was again videotaped to see if the dynamics went back to normal. From the videos, we then extracted distances between all fish and built social networks through a Markov chain approach (see [6]). We carried out 3 replicates for the cave mollies and 2 replicates for the surface mollies. In the following we will use the following abbreviations for the 3 treatments that were performed with each group: “G12a” for the first treatment with 12 fish, “G6” for the treatment with 6 fish, and “G12b” for the second treatment with 12 fish. We are aware that our treatments altered both number of individuals per group as well as individuals per area in the observational arena. However, for reasons of clarity we will refer to the change in individual numbers in the arena (G12a to G6 to G12b) as density change.

#### Individual tagging and housing of experimental fish before the recordings

Upon capture, fish were tagged immediately on site using VIE colours (see [38] for similar approach in the laboratory). To do so, fish were anesthetized with clove oil and carefully injected with VIE colours at different locations along their dorsal surface. VIE colours are visible in both day-light and infra-red light (as bright white spots) and this allowed us to individually recognize our fish in both environments. After the tagging, we transferred 12 individuals (6 males, 6 females) into a cubic net cage (60 cm by 60 cm by 60 cm; mesh-width 2 mm, Fig 1d). This cage was placed in a shallow area at both sites and the net bottom was covered entirely with natural gravel found around the cage. Water depth in the cage was about 5 cm and we ensured constant but slow water through flow. Afterwards, we left the fish within the cage overnight.

#### Video recording of social networks

The next morning, we filmed the fish with a full HD camera (Canon XF200, recording at 50 fps) which is able to record both in infrared light (used in the cave with an additional IR light, Dedolight Redzilla DLOBML-IR860, 860 nm) and normal light (used outside the cave in the El Azufre river). The camera was fixed centrally above the cage with a custom made tripod (Fig. 1d). While water through-flow was allowed during the night before the tests to enable food items reaching the cage, we closed any inflow into the cage by covering the outside walls with plastic foil during the experimental recordings. This was necessary to limit surface reflections due to water movements which would have hampered proper tracking of the fish. To handle fish, we used small head lights that were covered with red foil to minimize disturbances through direct light (see [39] for a similar approach when rearing cave fish in the laboratory). Once recordings were started, all experimenters left the site to avoid disturbances. After recording 12 fish for 45 min (G12a), we haphazardly selected and removed 3 males and 3 females from the cage. These fish were put into a similar net cage ca. 5 meters downstream. This procedure reduced the number of fish inside the cage by 50% (G6) and was inspired by experiments on Trinidadian guppies in natural ponds [2]. We recorded the cage for another 45 minutes as described before and then reintroduced the 6 previously removed fish back into the cage and recorded for a last period of 45 minutes (G12b). We measured body size (SL) of all experimental fish to the nearest millimetre. The body lengths of cave and surface mollies did not differ (mean body length of cave mollies = 37 mm ± 1.1 SE, mean body length of surface mollies = 38 mm ± 1.4 SE, *p* = 0.7 in a two samples *t*-test).

### Modelling of social dynamics

From all video recordings (3 videos per group, G12a, G6 and G12b), we analysed footage of 3 minutes which started 35 min after the recording was initialized. Pre-trials found fish in the cage to behave naturally ca. 10 min after introduction and by conservatively extending this period to 35 min we made sure to have undisturbed social interactions. We analysed only 3 minutes of each video after this necessary acclimation period (35 min) and not the full remaining 10 min (that add up to 45 min) mainly for two reasons: (i) The experimenters left the site after the camera recording was started and had to come back to stop the recording. In the cave it took about 5 min to reach the experimental site and torches and head lamps necessary to safely walk inside the cave might have illuminated the chamber where the experimental cage was placed during this 5 min. Therefore, to ensure that recordings were done without any light disturbance, it was necessary to discard the last 7 min. (ii) Tracking of the individual fish was time consuming for recordings both at the surface and the cave because of the low contrast between fish and background. We used natural gravel (greyish to blackish in coloration) found at each site and the water was milky due to oxidation of H_2_S. Therefore, tracking 3 min from each video was the reasonable maximum under these difficult field conditions. For those 3 min of footage, we individually tracked the positions of all fish using the video tracking software Ethovision XT10. As mentioned before, automated detection and tracking of individual fish was difficult due to the low contrast in the videos and tracks had to be corrected manually frame by frame. Although videos were recorded at 50 fps, we sampled fish positions at a constant rate of 5 fps. The resulting X-Y-position data were then exported and analysed further with custom made software.

#### The Markovian Model

As a basis for the investigation of the social dynamics of the mollies we used the stochastic model of Wilson et al. [6]. It can explain a number of aspects of social dynamics, including the percentage of time the individuals are social (or alone), the number and lengths of contact phases between the individuals, and the number and lengths of phases of individuals being alone. The model can be characterized by specifying the 3 probabilities of leaving the current nearest neighbour (*P*_leave_nn_), of discontinuing social contact in general (*P*_s→a_), and of discontinuing being alone (*P*_a→s_). The reciprocal values of theses probabilities are proportional to the mean lengths of contact phases with the same neighbour, of phases of being social (with any neighbour), and of phases of being alone, respectively. In addition, the probability of switching social partners within the period of social time (*P*_switch_nn_) can be derived from the model (see S1 Text for details on the model).

The data points for the construction of this model were collected by observations of focal individuals. Every *t* seconds the nearest neighbour of the focal individual was recorded, where two individuals were regarded as being neighbours of each other, if their distance was smaller than a value d. If no conspecific was within a radius of d, the individual was regarded as being alone. For our study we chose *t* = 5 seconds and *d* = 8 cm (see S1 Text for an explanation of this choice).

The four model probabilities are estimated as simple proportions (see S1 Text for more details). We compared them by looking at their 95 % confidence intervals using the function binom.test in R [40]. In our study, we have to be careful, because all focal individuals were observed simultaneously. This means, each contact phase was observed twice (once for each individual as a focal individual). To make sure that the confidence intervals of the probability *P*_leave_nn_ (of discontinuing g a contact phase) are not biased, we divided the numbers of data points used for their computation by 2. The confidence intervals of the probabilities P_s→a_ (of discontinuing a phase of being social) and *P*_switch_nn_ (of switching social partners within the period of social time) will also be affected by the simultaneous observation, although less strongly. Here, we reported confidence intervals for both the observed numbers of data points and the numbers of data points divided by 2. The correct value will be somewhere in between.

#### Influence of density changes

When the fish density changes, as is the case in our treatments through the change in numbers of individual fish in the arena, we should observe changes in the social dynamics, unless the fish work actively against them. For example, if the density decreases, the encounter probabilities between fish should also decrease. As a consequence, the lengths of phases of being alone should increase. In terms of the model probabilities this means that with decreasing density *P*_a→s_ also decreases. At the same time, the lengths of social phases decrease because after leaving a neighbour there will be a higher chance of being alone, i.e., with decreasing density *P*_s→a_ increases. As a result, the overall time of being social should decrease with decreasing density. The length of contact phases with the same neighbour should remain constant, i.e. *P*_leave_nn_ should not change. Consequently, with decreasing social time the number of switches of social partners should also decrease, i.e. *P*_switch_nn_ decreases.

To quantify the expected effects of changes in density on social dynamics, we performed a simulation of random movements of individuals, which has already been used by Wilson et al.[6]. A detailed description can be found in S1 Text.

#### Randomisation tests

As most measures taken from the behaviour of individuals in a group are dependent, we used randomisation tests to compare individual behaviour between the treatments. In each randomisation step the values of each individual were swapped between the treatments with a probability of 0.5. In other words, the observed values for each individual were randomly assigned to the treatments. For each test we performed 10^5^ randomisation steps.

## Results

Surface mollies showed lower probabilities of leaving the current social partner and of leaving any social partner than cave mollies (Fig 2). For G12 this was also the case for the probability of switching between social partners. The similarity of the switching probabilities for G6 is probably coincidental and only caused by cave mollies not actively working against density changes (see next paragraph). Surface mollies also showed a higher probability to join a new social partner once they were alone compared to cave mollies (but this difference was less pronounced for G12 than for G6, Fig 2).

**Fig 2.**
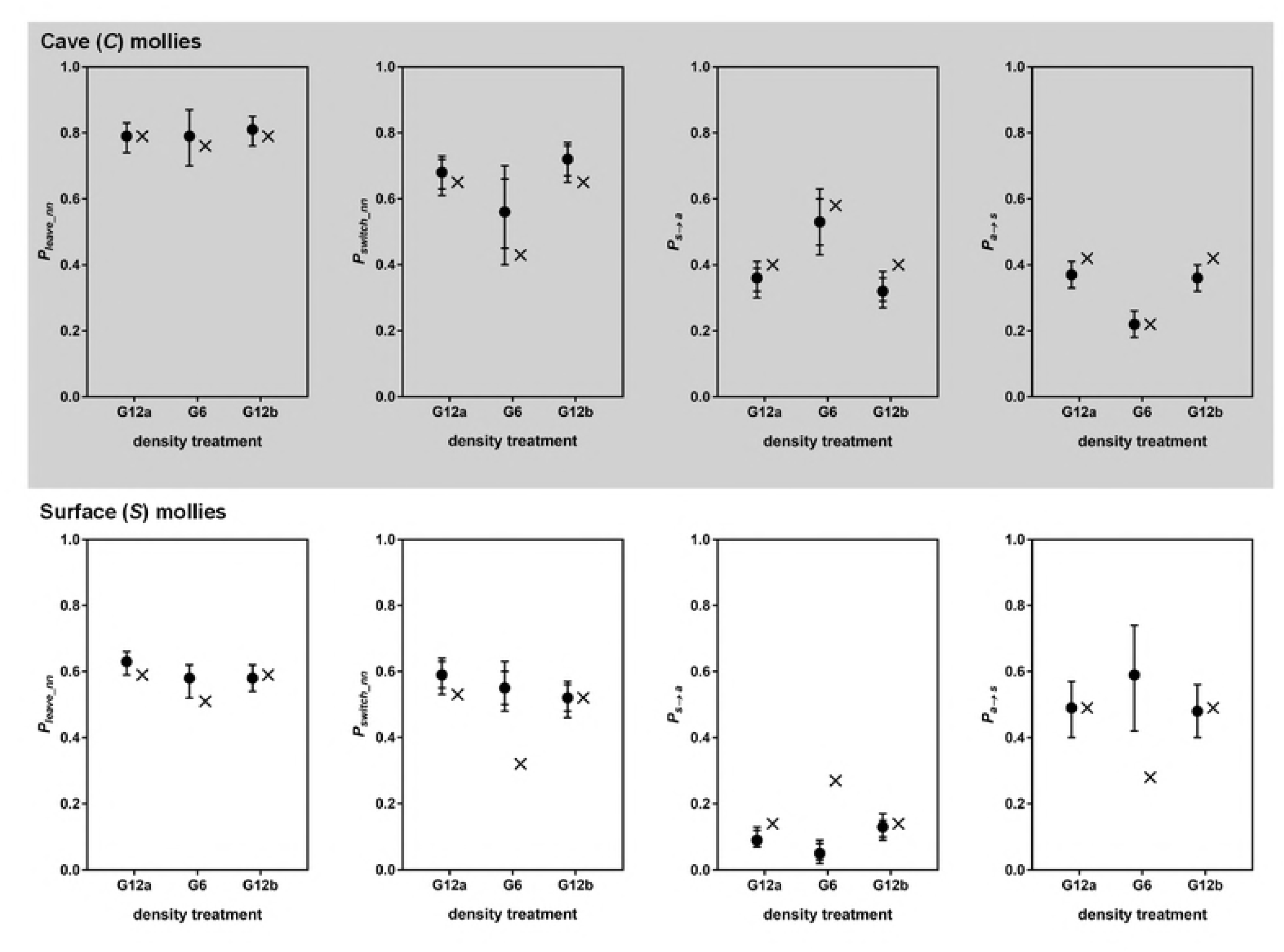
Estimated model probabilities (plus 95 % confidence intervals) for cave (a-d) and surface mollies (e-h). For the probabilities *P*_switch_nn_ and *P*_s→a_ two confidence intervals are shown for the reasons explained in the methods. The intervals with the longer horizontal lines were computed based on half of the data points. The probabilities resulting from a random walk simulation are marked by x’s. The parameters of the simulation were chosen such that it roughly reproduced the observed model probabilities of the groups of 12 fish.

### Effect of density changes on cave molly social dynamics

The social dynamics of cave mollies changed with changing density (from G12a to G6) as predicted by the random walk simulation. The probabilities *P*_a→s_ and *P*_switch_nn_ decreased while *P*_s→a_ increased, when the density decreased, i.e. the lengths of phases of being alone increased and the lengths of social phases decreased. The confidence intervals of *P*_s→a_ for the two treatments with 12 fish (G12a and G12b) were very similar but do not overlap with the confidence interval for the 6 fish treatment (G6). The same holds for of *P*_a→s_. Also, as predicted *P*_leave_nn_ did not seem to be influenced by density changes, i.e. the length of contact phases with the same social partner did not differ between G12 and G6.

The simulation predicted not only the trend of changes in social dynamics, but also approximately its magnitude. We chose the simulation parameters such that the simulation roughly reproduced the social dynamics of the treatments with 12 fish and then ran the simulation with 6 fish removed. It then generally reproduced the social dynamics that we observed in the treatment with 6 fish (Fig 2a-d). The percentage of overall time spent being social dropped from 52% (51% in the simulation) for the groups of 12 fish to 29% (27% in the simulation) for groups of 6 fish. This suggests that cave mollies followed the same individual behaviour pattern and did not adapt their behaviour to changes in density.

### Effect of density changes on surface molly social dynamics

For surface mollies the results were markedly different. The confidence intervals of all model probabilities overlapped between the treatments with 12 and with 6 fish (Fig 2e-h). Also, the observed probabilities in the treatment with 6 fish differed considerably from the predictions of the simulation (Fig 2e-h). The percentage of overall time spent being social increased from 81% (78% in the simulation) for the groups of 12 fish to 92% (50% in the simulation) for groups of 6 fish. This suggests that surface mollies worked actively against density changes.

In order to find out, which behavioural parameters surface mollies changed when the density decreased, we investigated three different aspects a) whether fish swam at higher speeds to maintain their social dynamics under lower densities, b) whether fish reduced area usage to a particular part of the net cage to compensate for lower densities and c) whether the polygon formed by the group decreased which might compensate for the density reduction (while allowing fish to visit most of the net cage).

### Swimming distances in cave and surface-dwelling mollies

In our analysis of individual swimming distances we included only those 6 individuals of each group that were present in all 3 treatments. To compare the swimming distances between G12 and G6 treatments, we computed the difference between individual total swimming distances in each treatment. The mean of these differences over all individuals constituted our test statistic.

Swimming distances of cave mollies did not change with density (Tables 1 and 2) which is in line with the results regarding their social dynamics, which suggested that cave mollies did not change their behaviour. Surface mollies, however, on average swam longer distances in groups of 12 than in groups of 6, although only the difference between G12a and G6 is significant (Tables 1 and 2). This result shows that the maintenance of sociality of surface mollies in the G6 treatment could not be explained by greater swimming distance (i.e. higher swimming speeds) in G6.

**Table 1.**
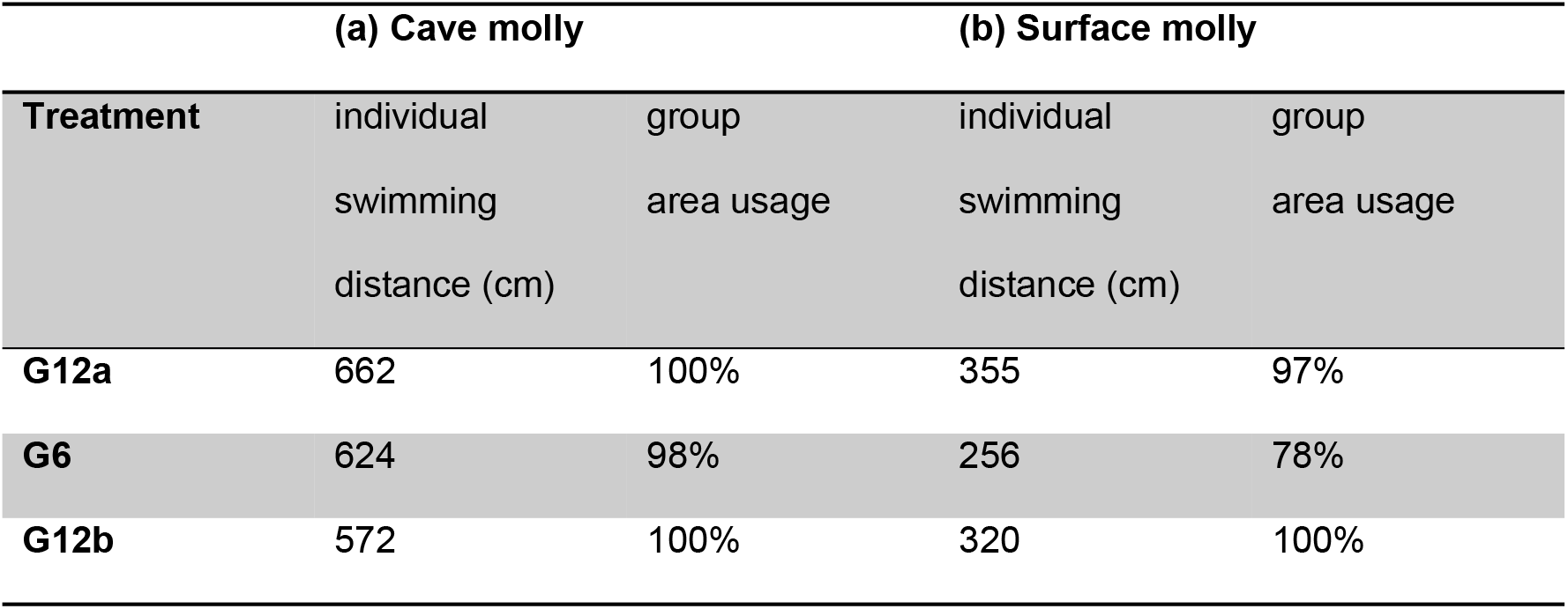
Mean individual swimming distances of the 6 fish that were present in all 3 treatments and mean area usage of the complete groups of the cave (a) and the surface mollies (b). The area usage indicates the percentage of squares visited in a 4×4 grid.

|  | **(a) Cave molly** |  | **(b) Surface molly** |  |
| --- | --- | --- | --- | --- |
| **Treatment** | individual | group | individual | group |
|  | swimming | area usage | swimming | area usage |
|  | distance (cm) |  | distance (cm) |  |
| **G12a** | 662 | 100% | 355 | 97% |
| **G6** | 624 | 98% | 256 | 78% |
| **G12b** | 572 | 100% | 320 | 100% |

**Table 2.**
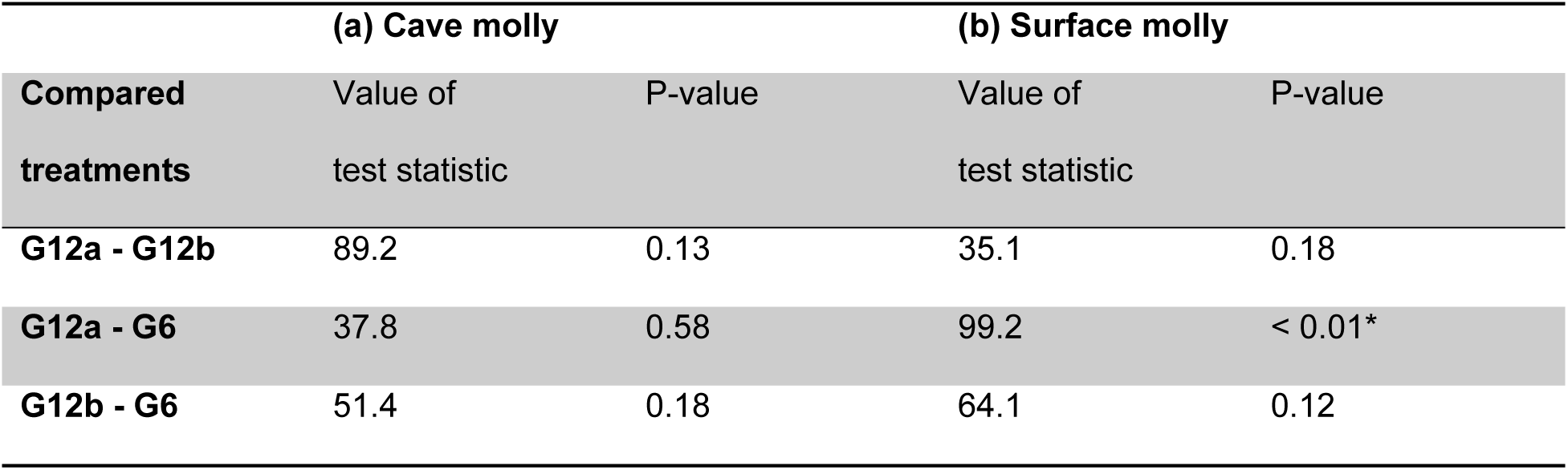
Results of randomisation tests (N=10^5^ repetitions) regarding differences in individual swimming distances for the cave (a) and the surface mollies (b). Significant results are marked with a star.

### Area usage in cave and surface-dwelling mollies

To measure the group area use, we divided the total area (60 × 60 cm^2^) into 16 squares of 15 × 15 cm^2^ and counted the number of squares visited by some individual of the group. The group area usage of cave mollies did not change with density (Table 1), which again suggests that cave mollies did not change their behaviour. Surface mollies, however, reduced their group area usage on average by 21% when the number of fish decreased. Using our random walk simulation we tested whether this reduction would be sufficient to maintain the observed probabilities of social dynamics for G6 and thus account for the density compensation. Our simulation indicated that this was not the case (*P*_s→a_ observed=0.05, simulated with area reduction=0.24, simulated without area reduction=0.27; *P*_a→s_ observed = 0.59, simulated with = 0.33 and without area reduction = 0.28; overall social time observed = 92% and simulated with = 58% and without area reduction = 50%). For surface mollies, an area reduction of 60% (instead of only 21%) would be necessary to achieve the observed probabilities of social dynamics for G6.

### Size of the polygons formed by the groups of cave and surface-dwelling mollies

If surface mollies were able to reduce the size of a convex polygon formed by the group as density decreased, then this could allow them to largely maintain their social dynamics while still being able to visit large parts of the available area as a group. In order to investigate this issue we computed the mean perimeter and mean area of the convex polygons formed by both cave and surface mollies in the three treatments. In contrast to the analyses of individual measures, we took the complete groups into account and not just the 6 individuals from the G6 treatments. However, we had to take into account that the size of the polygon decreases as density decreases, even if the fish did not change their behaviour. To estimate the magnitude of this effect, we computed the mean perimeter and mean area from the random walk simulations used for the analysis of the social dynamics.

The behaviour of cave mollies was again predicted by the simulation (Table 3). The reductions of both the mean perimeter and the mean area of the convex polygons formed by the group could be explained by a reduction in density, without individuals changing their behaviour.

**Table 3.** Mean perimeter and mean area of the convex polygons formed by the groups in the three treatments of the cave (a) and the surface mollies (b). For the groups of size 6 the reduction relative to the values of the groups of size 12 is shown. The simulation results are the mean results from 1000 random walks. The values marked with an asterisk were obtained from a simulation with the movement rules explained in Fig 3.

|  | Mean<br>perimeter<br>[cm] | %reduced |  | Mean area<br>[cm<sup>2</sup>] | %reduced |  |
| --- | --- | --- | --- | --- | --- | --- |
|  | observed | observed | simulated | observed | observed | simulated |
| **(a) Cave molly** |  |  |  |  |  |  |
| **G12a** | 187 |  |  | 2172 |  |  |
| **G6** | 157 | 15 | 17 | 1285 | 38 | 42 |
| **G12b** | 181 |  |  | 1977 |  |  |
| **(b) Surface molly** |  |  |  |  |  |  |
| **G12a** | 103 |  |  | 620 |  |  |
| G6 | 49 | 55 | 17 (39)* | 93 | 86 | 42 (70)* |
| G12b | 112 |  |  | 698 |  |  |

As expected, the results were different for the surface mollies (Table 3). Here, the reductions of both the mean perimeter and the mean area of the polygons were much larger than the density reduction alone can explain. In order to test the significance of these reductions we determined the 0.025 percentiles of the distribution of mean perimeters and mean areas using 10^4^ repetitions of our random walk simulation. For both measures the observed values (55% reduction of the perimeter and 86% reduction of the area) were greater than the 0.025 percentiles (25% reduction of the perimeter and 56% reduction of the area). Therefore, the size reduction of the polygons can be regarded as significant.

We explored individual-based movement rules that fish could perform to achieve changes in polygon size as a result of changes in density. A simple movement rule which substantially changed polygon size while maintaining the social dynamics requires fish to move back into the polygon at the next time step in the simulation when they are at the vertex of a convex polygon and without a neighbour within two body lengths (Fig 3, Table 3b).

**Fig 3.**
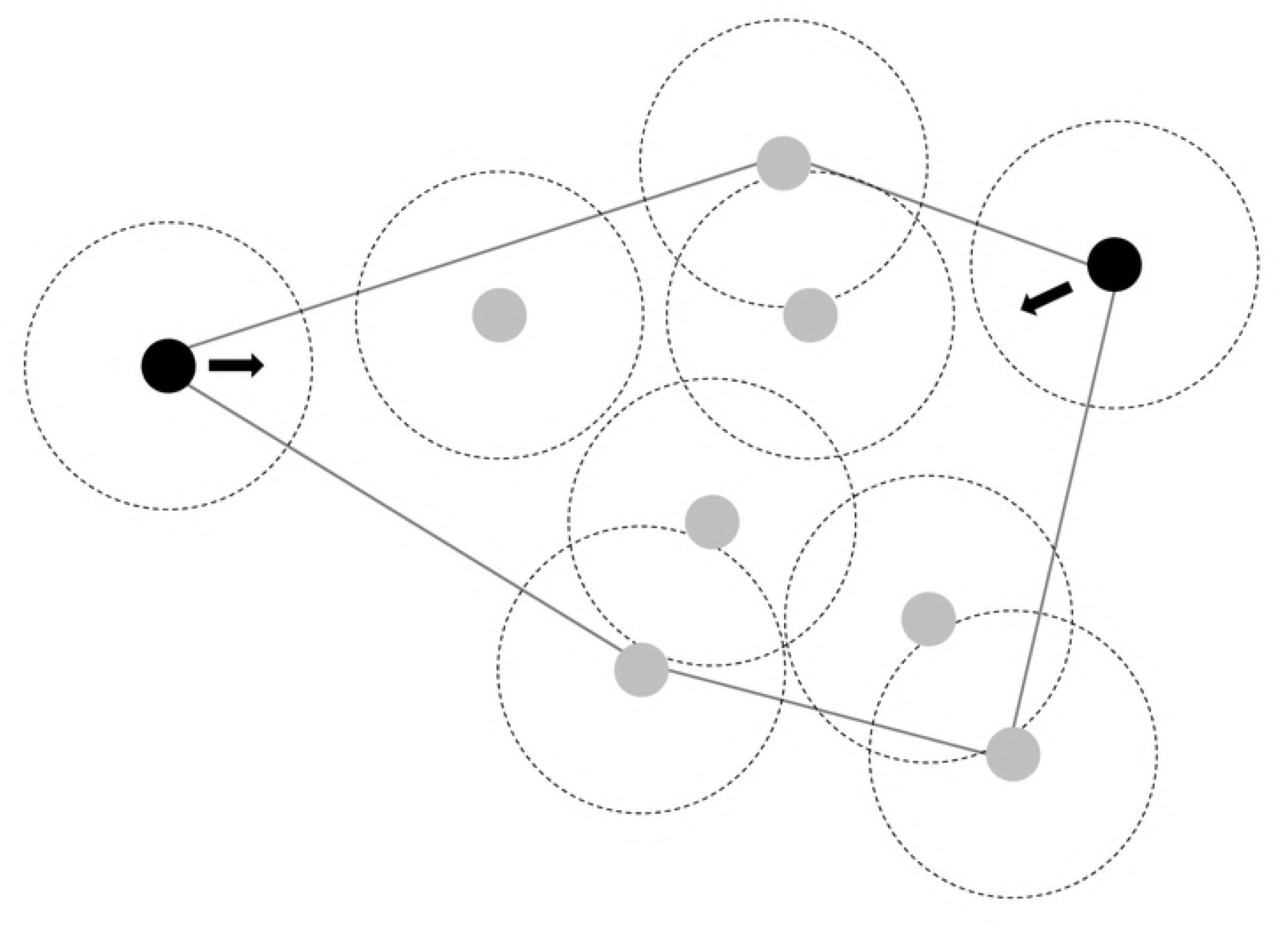
Illustration of individual movement rules of fish which could largely explain observed changes in polygon size (in response to density changes) and maintain social dynamics. Individuals at the vertex of a polygon without a neighbour within 8 cm move at the next time step back into the polygon area (instead of further out).

### Testing alternative explanations for reduction in polygon size in surface-dwelling mollies

The observed reduction in polygon size in surface mollies could be result of a fundamental change in behaviour from a fission-fusion system to schooling when density changed. To investigate this issue we measured the polarization of the 6 fish that were present in all three treatments by computing the sum of the unit velocity vectors of these 6 individuals every 3 seconds. Then we computed the mean of these sums. The interval of 3 s was chosen, because sometimes fish did not move for almost 3 s, leading to an undefined unit vector. The sum of the vectors is always in the range 0 – 1. The value 1 occurs, if all fish swim in exactly the same direction, the value 0 occurs, if all swimming directions cancel each other out. Surface mollies generally appeared to have a higher polarization than cave mollies but did not change their degree of polarization when the density decreased (Table 4).

**Table 4.** Mean sums of unit velocity vectors of the 6 fish present in all 3 treatments of the cave and the surface mollies.

|  | Cave mollies | Surface mollies |
| --- | --- | --- |
| Treatment | Mean sum of unit vectors |  |
| G12a | 0.37 | 0.48 |
| G6 | 0.36 | 0.48 |
| G12b | 0.33 | 0.45 |

## Discussion

Overall, surface mollies spent much more time being social than cave mollies. As predicted, surface mollies were less likely to leave a current social partner and to stop being social than cave mollies. There was also a trend for less partner switching in surface mollies for groups of 12 individuals compared to cave mollies. Given the unusually high bird predation outside the cave, moving away from partners and, to a lesser extent switching partners is presumably risky and should promote the formation of stronger social bonds between individuals. This is an aspect of our research that deserves further attention and has also been highlighted in comparisons of the social structure of high- and low-predation populations in guppies, *P*. *reticulata* [7]. Cave and surface mollies also differed in the probability to become social after a phase of being alone (*P*_a→s_). The low probability to stop being social and the high probability to re-join a conspecific once alone combined to produce a high percentage of time spent social (81–92%) in surface mollies which is comparable to the most extreme high-predation populations of guppies (*P. reticulata*) in Trinidad [41]. In contrast the social dynamics of the cave mollies largely followed a null-model (with no social interactions). Magurran et al. ([42]) reported that in a high-predation guppy-population (*P. reticulata*) shoaling was significantly reduced (by 2–2.5 times in males and females, respectively) after the fish had been introduced to a low-predation area and left there for over 50 years to reproduce and adapt. By comparison shoaling behaviour seems to have disappeared completely in our cave mollies and it is estimated that ancestral surface mollies entered this cave approximately 100 ka ago [10]. The consequences of this complete lack of shoaling behaviour (despite the very high densities of fish, in some chamber more than 200 individuals per square metre [43]) for the foraging ecology and mating behaviour are largely unknown and present interesting opportunities for future research. It is known from lab-reared populations of cave mollies that they show shoaling behaviour under daylight conditions and are only asocial under low-light conditions (in contrast to surface mollies which maintain some shoaling behaviour under low-light conditions too, [11]).

The surface mollies displayed an ability to maintain their social dynamics despite density changes which was not observed in cave mollies. It is likely that these substantial differences in the social dynamics between cave and surface mollies are the evolutionary outcome to an environment with markedly different predatory threats [14, 15]. The cave habitat harbours fish-eating bugs (*Belostoma* sp. [44, 45]), crabs [46] as well as spiders [47] – all of which sit at the waters’ edge above the water level and hunt by holding mandibles or legs into the water waiting for a cave molly to swim near and eventually touch it. Thus, their hunting style is non-visual. However, anti-predator benefits of being social are proposed mostly to reduce risks posed by visually hunting predators [1]. Following this logic, cave mollies face no strong selection pressure to remain social in their cave environment. In contrast, the surface mollies are subject to very high predation pressures by fish-eating birds [20]. This is due to the fact that the sulfidic water is almost free of dissolved oxygen and forces fish to the surface in order to perform aquatic surface respiration [48]. For cave mollies that live in the very same sulfidic waters this has no predation-related consequences (there are no birds hunting in the cave) but in the surface mollies this makes them highly accessible to birds such as herons and kingfishers and bird predation rates were found to be increased 20-fold in sulfidic waters compared to other clear-water surface streams[20].

The fact that the surface mollies, just like guppies [2], compensated extreme density changes to maintain their social dynamics suggests that this particular level of dynamics possesses an important adaptive value which is presumably linked to anti-predator benefits and information exchange. So, with half the number of fish in the arena, how can surface mollies still keep their social dynamic (e.g., similar probabilities to switch from social to asocial and vice versa under all densities) while simultaneously visiting most of the available space in the arena? A reduction of the used area in the arena was detectable in the low density treatment (21% reduction), but a much greater reduction (about 60%) would have been necessary to explain the observed compensation in the social dynamics by the area usage hypothesis. The fact that the reduction in the polygon size spanned by the group was much greater than one would expect by a reduction in density alone suggests a rather simple mechanism for maintaining social dynamics under changing density contexts. Such a reduction in polygon size could be achieved by fish moving back into the group when they are at the vertex of a convex polygon spanned by their neighbours that are outside the social range (in our case outside an 8 cm radius, see Fig 3). The theory of marginal predation [49] suggests that if predators attack the closest prey, then those on the edge of groups should experience greater risk [50, 51]. Thus, a strong selection pressure for individuals to move back into the group, e.g. swim within the vertex polygon, is likely to be common in group-living animals (see [52]). Furthermore, a possible mechanism that enables surface mollies to maintain their social dynamics under varying densities is known in starlings, *Sturnus vulgaris*, that take their six to seven nearest neighbours into account when flying in large flocks [53], monitoring the nearest neighbours and their relative distance and spatial position.

We did not find any indication that the surface mollies switched from fission-fusion behaviour to schooling behaviour in response to being at a lower density (Table 4). Also, the polarization of surface mollies was much lower than for typical schooling behaviour observed in other poeciliid studies (showing values of 0.75 and higher, see [54]). Therefore we conclude that increased polarization was not responsible for the observed reduction in the polygon size in surface mollies.

It is a particular strength of the Markov model approach in combination with the density manipulations that we can provide insights into selection processes that potentially operated on different components of the social behaviour of these molly populations. It is a weakness of our present study, however, that we could make this comparison for only one population inside and one outside a cave (in the absence of other available population pairs inside and outside caves in that region). Further research on other populations that inhabit environments both inside and outside caves will be necessary to complete the picture, for example using the Mexican cave tetra (*Astyanax mexicanus*) and its surface living relatives [55, 56]. At least for the density compensation similar observations on other fish species already exist (see [2]).

### Ethics

Experiments reported in this study were carried out in accordance with the recommendations of “Guidelines for the treatment of animals in behavioural research and teaching” (published in Animal Behaviour 1997) and under the authorization of the Mexican government (DGOPA.09004.041111.3088, PRMN/DGOPA-003/2014, PRMN/DGOPA-009/2015, and PRMN/DGOPA-012/2017, issued by SAGARPA-CONAPESCA-DGOPA).

### Data accessibility

Our data will be deposited at Dryad upon acceptance for publication.

### Authors’ contributions

DB, SK, PR and JK designed the study, DB, LA-R and JK performed the experiments, SK, DB, PR, JL and JK analysed the data, DB, SK and JK wrote the first draft of the manuscript. All authors worked on the manuscript and approved the final version.

### Funding

We received funding by the DFG (BI 1828/2–1 to DB and, RO 4766/2–1 to PR) as well as the IGB seed money program (to JK) and the Leibniz-Competition (SAW-2013-IGB-2 to JK).

## Acknowledgements

This experiment was inspired by H.G. Wells’ “The Time Machine” in which Wells described the adaptations of surface-dwelling and subterranean populations of humans – the Eloi and the Morlocks, respectively. We like to thank the people at Teapa and Tapijulapa for their great hospitality during our various field trips to the cave and surrounding waters. Furthermore, we like to thank Jonas Jourdan, L. Alex Jordan, Carolin Sommer-Trembo as well as Claudia Zimmer for their help in the field.

## Supporting information

**S1 Text: Details on the Markov Chain Method for analysing social dynamics**.

## References

1. Krause J, Ruxton GD. Living in groups. Oxford: Oxford University Press; 2002.

2. Wilson ADM, Krause S, Ramnarine IW, Borner KK, Clément RJG, Kurvers RHJM, et al. Social networks in changing environments. Behavioral Ecology and Sociobiology. 2015;69(10):1617–29. doi: 10.1007/s00265-015-1973-2.

3. Wilson ADM, Brownscombe JW, Krause J, Krause S, Gutowsky LFG, Brooks EJ, et al. Integrating network analysis, sensor tags, and observation to understand shark ecology and behavior. Behavioral Ecology. 2015;26(6):1577–86. doi: 10.1093/beheco/arv115.

4. Deacon AE, Jones FA, Magurran AE. Gradients in predation risk in a tropical river system. Current Zoology. 2018:zoy004-zoy. doi: 10.1093/cz/zoy004.

5. Magurran AE. Evolutionary ecology: The Trinidadian guppy Oxford: Oxford University Press; 2005.

6. Wilson AM, Krause S, James R, Croft D, Ramnarine I, Borner K, et al. Dynamic social networks in guppies (*Poecilia reticulata*). Behavioral Ecology and Sociobiology. 2014:1–11. doi: 10.1007/s00265-014-1704-0.

7. Kelley JL, Morrell LJ, Inskip C, Krause J, Croft DP. Predation Risk Shapes Social Networks in Fission-Fusion Populations. PLoS ONE. 2011;6(8):e24280. doi: 10.1371/journal.pone.0024280.

8. Tobler M, Kelley JL, Plath M, Riesch R. Extreme environments and the origins of biodiversity: adaptation and speciation in sulfide spring fishes. Molecular Ecology. 2018:n/a-n/a. doi: 10.1111/mec.14497.

9. Riesch R, Tobler M, Plath M. Hydrogen Sulfide-Toxic Habitats. In: Riesch R, Tobler M, Plath M, editors. Extremophile Fishes: Springer International Publishing Switzerland 2015; 2015.

10. Pfenninger M, Lerp H, Tobler M, Passow C, Kelley JL, Funke E, et al. Parallel evolution of cox genes in H2S-tolerant fish as key adaptation to a toxic environment. Nature Communications. 2014;5:3873. doi: 10.1038/ncomms4873 https://www.nature.com/articles/ncomms4873#supplementary-information.

11. Bierbach D, Lukas J, Bergmann A, Elsner K, Höhne L, Weber C, et al. Insights into the Social Behavior of Surface and Cave-Dwelling Fish (*Poecilia mexicana*) in Light and Darkness through the Use of a Biomimetic Robot. Frontiers in Robotics and AI. 2018;5(3). doi: 10.3389/frobt.2018.00003.

12. Riesch R, Schlupp I, Tobler M, Plath M. Reduction of the association preference for conspecifics in cave-dwelling Atlantic mollies, Poecilia mexicana. Behavioral Ecology and Sociobiology. 2006;60(6):794–802. doi: 10.1007/s00265-006-0223-z.

13. Plath M, Schlupp I. Parallel evolution leads to reduced shoaling behavior in two cave dwelling populations of Atlantic mollies (*Poecilia mexicana*, Poeciliidae, Teleostei). Environmental Biology of Fishes. 2008;82(3):289–97. doi: 10.1007/s10641-007-9291-9.

14. Tobler M, Schlupp I, Heubel KU, Riesch R, de Leon FJ, Giere O, et al. Life on the edge: hydrogen sulfide and the fish communities of a Mexican cave and surrounding waters. Extremophiles. 2006;10(6):577–85. PubMed PMID: 16788733.

15. Greenway R, Arias-Rodriguez L, Diaz P, Tobler M. Patterns of Macroinvertebrate and Fish Diversity in Freshwater Sulphide Springs. Diversity. 2014;6(3):597. PubMed PMID: doi:10.3390/d6030597.

16. Bierbach D, Schulte M, Herrmann N, Zimmer C, Arias-Rodriguez L, Indy JR, et al. Predator avoidance in extremophile fish. Life. 2013;3(1):161–80. PubMed PMID: doi:10.3390/life3010161.

17. Borner K, Krause S, Mehner T, Uusi-Heikkilä S, Ramnarine I, Krause J. Turbidity affects social dynamics in Trinidadian guppies. Behavioral Ecology and Sociobiology. 2015:1–7. doi: 10.1007/s00265-015-1875-3.

18. Croft D, Krause J, Darden S, Ramnarine I, Faria J, James R. Behavioural trait assortment in a social network: patterns and implications. Behavioral Ecology and Sociobiology. 2009;63(10):1495–503. doi: 10.1007/s00265-009-0802-x.

19. Croft DP, James R, Thomas POR, Hathaway C, Mawdsley D, Laland KN, et al. Social structure and co-operative interactions in a wild population of guppies (*Poecilia reticulata*). Behavioral Ecology and Sociobiology. 2006;59(5):644–50. doi: 10.1007/s00265-005-0091-y.

20. Riesch R, Oranth A, Dzienko J, Karau N, SchießL A, Stadler S, et al. Extreme habitats are not refuges: poeciliids suffer from increased aerial predation risk in sulphidic southern Mexican habitats. Biological Journal of the Linnean Society. 2010;101(2):417–26. doi: 10.1111/j.1095-8312.2010.01522.x.

21. Parzefall J. A review of morphological and behavioural changes in the cave molly, *Poecilia mexicana*, from Tabasco, Mexico. Environmental Biology of Fishes. 2001;62(1–3):263–75.

22. Gordon MS, Rosen DE. A cavernicolous form of the poeciliid dish *Poecilia sphenops* from Tabasco, México. Copeia. 1962;1962:360–8.

23. Tobler M, Riesch R, García de León FJ, Schlupp I, Plath M. A new and morphologically distinct population of cavernicolous Poecilia mexicana (Poeciliidae: Teleostei). Environmental Biology of Fishes. 2008;82(1):101–8. doi: 10.1007/s10641-007-9258-x.

24. Miller RR. Freshwater fishes of Mexico. Chicago: Chicago University Press; 2006.

25. Greven H. Gonads, genitals, and reproductive biology. In: Evans JP, Pilastro A, Schlupp I, editors. Ecology and evolution of poeciliid fishes. Chicago: Chicago University Press; 2011. p. 57–87.

26. Parzefall J. Zur vergleichenden Ethologie verschiedener *Mollienesia*-Arten einschließlich einer Höhlenform von Mollienesia sphenops. Behaviour. 1969;33(1):1–38. PubMed PMID: 5815891.

27. Niemeitz A, Kreutzfeldt R, Schartl M, Parzefall J, Schlupp I. Male mating behaviour of a molly, *Poecilia latipunctata*: a third host for the sperm-dependent Amazon molly, *Poecilia formosa*. Acta ethologica. 2002;5(1):45–9. doi: 10.1007/s10211-002-0065-2.

28. Bierbach D, Makowicz AM, Schlupp I, Geupel H, Streit B, Plath M. Casanovas are liars: behavioral syndromes, sperm competition risk, and the evolution of deceptive male mating behavior in live-bearing fishes. F1000Research. 2013;2:75.

29. Bierbach D, Schulte M, Herrmann N, Tobler M, Stadler S, Jung CT, et al. Predator-induced changes of female mating preferences: innate and experiential effects. BMC Evolutionary Biology. 2011;11(1). doi:10.1186/1471-2148-11-190.

30. Plath M, Parzefall J, Schlupp I. The role of sexual harassment in cave and surface dwelling populations of the Atlantic molly, Poecilia mexicana (Poeciliidae, Teleostei). Behavioral Ecology and Sociobiology. 2003;54(3):303–9. doi: 10.1007/s00265-003-0625-0.

31. Bierbach D, Arias-Rodriguez L, Plath M. Intrasexual competition enhances reproductive isolation between locally adapted populations. Current Zoology. 2017:zox071-zox. doi: 10.1093/cz/zox071.

32. Bierbach D, Klein M, Sassmannshausen V, Schlupp I, Riesch R, Parzefall J, et al. Divergent evolution of male aggressive behaviour: another reproductive isolation mechanism in extremophile poeciliid fishes. International Journal of Evolutionary Biology. 2012;2012(ID 148745):148745. doi: 10.1155/2012/148745.

33. Parzefall J. Rückbildung aggressiver Verhaltensweisen bei einer Höhlenform von Poecilia sphenops (Pisces, Poeciliidae). Zeitschrift für Tierpsychologie. 1974;35(1):66–84. doi: 10.1111/j.1439-0310.1974.tb00433.x.

34. Eifert C, Farnworth M, Schulz-Mirbach T, Riesch R, Bierbach D, Klaus S, et al. Brain size variation in extremophile fish: local adaptation versus phenotypic plasticity. Journal of Zoology. 2014:n/a-n/a. doi: 10.1111/jzo.12190.

35. Tobler M, Coleman SW, Perkins BD, Rosenthal GG. Reduced opsin gene expression in a cave-dwelling fish. Biology Letters. 2010;6(1):98–101. doi: 10.1098/rsbl.2009.0549.

36. Fontanier ME, Tobler M. A morphological gradient revisited: cave mollies vary not only in eye size. Environmental Biology of Fishes. 2009;86(2):285–92. doi: 10.1007/s10641-009-9522-3.

37. Körner KE, Schlupp I, Plath M, Loew ER. Spectral sensitivity of mollies: comparing surface-and cave-dwelling Atlantic mollies, Poecilia mexicana. Journal of Fish Biology. 2006;69(1):54–65. doi: 10.1111/j.1095-8649.2006.01056.x.

38. Bierbach D, Oster S, Jourdan J, Arias-Rodriguez L, Krause J, Wilson AM, et al. Social network analysis resolves temporal dynamics of male dominance relationships. Behavioral Ecology and Sociobiology. 2014:1–11. doi: 10.1007/s00265-014-1706-y.

39. Riesch R, Reznick DN, Plath M, Schlupp I. Sex-specific local life-history adaptation in surface-and cave-dwelling Atlantic mollies (*Poecilia mexicana*). Scientific Reports. 2016;6:22968. doi: 10.1038/srep22968 http://www.nature.com/articles/srep22968#supplementary-information.

40. R_Core_Team. R: A Language and Environment for Statistical Computing Vienna, Austria: R Foundation for Statistical Computing; 2013. Available from: http://www.Rproject.org/.

41. Magurran AE, Seghers BH. Variation in Schooling and Aggression Amongst Guppy (*Poecilia reticulata*) Populations in Trinidad. Behaviour. 1991;118(3):214–34. doi: https://doi.org/10.1163/156853991X00292.

42. Magurran AE, Seghers BH, Carvalho GR, Shaw PW. Behavioural consequences of an artificial introduction of guppies (*Poecilia reticulata*) in N. Trinidad: evidence for the evolution of anti-predator behaviour in the wild. Proceedings of the Royal Society of London Series B: Biological Sciences. 1992;248(1322):117–22. doi: 10.1098/rspb.1992.0050.

43. Jourdan J, Bierbach D, Riesch R, Schießl A, Wigh A, Arias-Rodriguez L, et al. Microhabitat use, population densities, and size distributions of sulfur cave-dwelling *Poecilia mexicana*. PeerJ. 2014;2:e490. doi: 10.7717/peerj.490.

44. Tobler M. Does a predatory insect contribute to the divergence between cave-and surface-adapted fish populations? Biology Letters. 2009;5(4):506–9. doi: 10.1098/rsbl.2009.0272.

45. Tobler M, Schlupp I, Plath M. Predation of a cave fish (*Poecilia mexicana*, Poeciliidae) by a giant water-bug (Belostoma, Belostomatidae) in a Mexican sulphur cave. Ecological Entomology. 2007;32(5):492–5. doi: 10.1111/j.1365-2311.2007.00892.x.

46. Klaus S, Plath M. Predation on a cave fish by the freshwater crab *Avotrichodactylus bidens* (Bott, 1969) (Brachyura, Trichodactylidae) in a Mexican sulfur cave. Crustaceana. 2011;84:411−8

47. Horstkotte J, Riesch R, Plath M, Jäger P. Predation on a cavefish (*Poecilia mexicana*) by three species of spiders in a Mexican sulfur cave. Bulletin of the British Arachnolocical Society. 2010;15:55–8.

48. Plath M, Tobler M, Riesch R, Garcia de Leon FJ, Giere O, Schlupp I. Survival in an extreme habitat: the roles of behaviour and energy limitation. Naturwissenschaften. 2007;94(12):991–6. PubMed PMID: 17639290.

49. Hamilton WD. Geometry for the selfish herd. Journal of Theoretical Biology. 1971;31(2):295–311. doi: https://doi.org/10.1016/0022–5193(71)90189–5.

50. Bumann D, Krause J, Rubenstein D. Mortality Risk of Spatial Positions in Animal Groups: The Danger of Being in the Front. Behaviour. 1997;134(13/14):1063–76.

51. Morrell LJ, Romey WL. Optimal individual positions within animal groups. Behavioral Ecology. 2008;19(4):909–19. doi: 10.1093/beheco/arn050.

52. Morrell LJ, Ruxton GD, James R. Spatial positioning in the selfish herd. Behavioral Ecology. 2011;22(1):16–22. doi: 10.1093/beheco/arq157.

53. Ballerini M, Cabibbo N, Candelier R, Cavagna A, Cisbani E, Giardina I, et al. Interaction ruling animal collective behavior depends on topological rather than metric distance: Evidence from a field study. Proceedings of the National Academy of Sciences. 2008;105(4):1232–7. doi: 10.1073/pnas.0711437105.

54. Herbert-Read JE, Krause S, Morrell LJ, Schaerf TM, Krause J, Ward AJW. The role of individuality in collective group movement. Proceedings of the Royal Society of London B: Biological Sciences. 2012;280(1752).

55. Protas M, Conrad M, Gross JB, Tabin C, Borowsky R. Regressive evolution in the Mexican cave tetra, *Astyanax mexicanus*. Current Biology. 2007;17(5):452–4. doi: http://dx.doi.org/10.1016/j.cub.2007.01.051.

56. Panaram K, Borowsky R. Gene-flow and genetic variability in cave and surface populations of the Mexican tetra, Astyanax mexicanus (Teleostei: Characidae). Copeia. 2005:409–16. doi:10.1643/CG-04-068R1.

